# Factorbook Motif Pipeline: A *de novo* motif discovery and filtering web server for ChIP-seq peaks

**DOI:** 10.1101/033670

**Authors:** Bong-Hyun Kim, Jiali Zhuang, Jie Wang, Zhiping Weng

## Abstract

**Summary:** High-throughput sequencing technologies such as ChIP-seq have deepened our understanding in many biological processes. *De novo* motif search is one of the key downstream computational analysis following the ChIP-seq experiments and several algorithms have been proposed for this purpose. However, most web-based systems do not perform independent filtering or enrichment analyses to ensure the quality of the discovered motifs. Here, we developed a web server Factorbook Motif Pipeline based on an algorithm used in analyzing ENCODE consortium ChIP-seq datasets. It performs comprehensive analysis on the set of peaks detected from a ChIP-seq experiments: (i) *de novo* motif discovery; (ii) independent composition and bias analyses and (iii) matching to the annotated motifs. The statistical tests employed in our pipeline provide a reliable measure of confidence as to how significant are the motifs reported in the discovery step.

**Availability:** Factorbook Motif Pipeline source code is accessible through the following URL. https://github.com/joshuabhk/factorbook-motif-pipeline

## 1 INTRODUCTION

Chromatin immunoprecipitation followed by sequencing (ChIP-seq) experiments have been crucial in understanding the regulation of many biological processes by revealing transcription factor (TF) binding sites *in vivo* on the whole genome scale (Solomon *et al*., 1988: Robertson *et al*., 2007). The information about TF binding provides unprecedented opportunities to understand the interactions and regulations of multiple TFs from many different cell or tissue types in a variety of conditions (Gerstein *et al*., 2012). Multiple lines of evidence favored the model in which TFs extensively interact with each other and perform their regulatory functions as components of larger multi-unit protein complexes over the model where a TF functions in isolation (Li *et al*., 2008; ENCODE Project Consortium *et al*., 2012). Moreover, TF binding frequently occurs without any discernible TF binding sites (or motif instances) through tethered binding (Wang *et al*., 2012){Wang:2013fp} or to hot-spots where multiple TFs binds simultaneously possibly in conjunction with the chromatin structures (Siersbæk *et al*., 2011). To understand the TF-DNA interactions, the underlying sequence features or motifs need to be analyzed. However, as shown in previous studies, up to a quarter of *de novo* discovered motifs appear to be random patterns (Kheradpour and Kellis, 2013). Currently available *de novo* motif discovery web tools (Machanick and Bailey, 2011) do not provide adequate quality control measures that can distinguish genuine motifs enriched in the ChIP-seq peaks from random DNA patterns. Here, we describe our *de novo* motif discovery web resource, named Factorbook Motif Pipeline, which provides an independent motif filtering step after *de novo* motif discovery in addition to comprehensive motif analyses.

## 2 METHODS AND IMPLEMENTATION

Factorbook Motif Pipeline performs *de novo* motif search, subsequent filtering and discovering similar motifs in the databases from genomic regions defined by user in NarrowPeak or BED formats sorted by peak strengths. The server uses 100bp around the peak summits of 500 peaks with strongest signals to *de novo* discover the motifs by MEME program as used in MEME-ChIP server (Bailey *et al*., 2009; Machanick and Bailey, 2011). The discovered motifs were statistically tested by (i) randomly selected genomic regions with the same GC contents matching to the peak regions and by (ii) flanking regions of the peaks. To test if the discovered motif is due to GC content bias, the server generates FIMO score (Grant *et al*., 2011) distribution by scanning the motif on the 300bp regions centered on the peak summit for the 500 peaks ranked from 500~1000. Then a control distribution is generated for the motif using FIMO on randomly selected GC-matched genomic regions excluding the peak regions. Empirical *P*-value for the actual distribution is calculated based on the mean and standard deviation from 100 random samplings. In addition, to test if the discovered motif is due to local sequences feature, the server uses flanking regions of the peaks ranked 500~1000, and calculates the enrichment of the motif measured by the number of peak regions that contain motif divided by the number of flanking regions that contain motif using FIMO. Details of the filtering method are described in our previous publication (Wang *et al*., 2012).

We carefully designed Factorbook Motif Pipeline so that our server can provide a template for future bioinformatics server development. We separated the frontend server that processes user requests and the backend server that computes the motif discovery, filtering and comparison. We implanted the server systems without relying on a database management system to reduce administrative burden and system overheads. It is achieved by introducing non-competing unique identifier and transferring the user submitted information and data file contained in a directory named after the unique identifier between the frontend and backend servers. We also used cherrypy (www.cherrypy.org), jinja2 (jinja.pocoo.org), google AJAX and google charts libraries that helped to improve code reusability and maintainability. We developed and tested our codes in python 2.7.3 and it will work with python 2.7 or above. The running time of the server depends on the number peaks in the user’s input file, and it typically takes couple hours to process since the server need to run computationally expensive MEME program to discover motifs and then need to run many random trials for the significance tests.

All codes and data can be found at the following URL. http://zlab.umassmed.edu/~kimb/factorbook_motif_pipeline/

## 3 EXAMPLE

We demonstrated the usage of Factorbook Motif Pipeline on two ChIP-seq peak datasets downloaded from ENCODE consortium. One dataset is the ChIP-seq for CTCF, a sequence-specific TF with well characterized motif; the other dataset is the ChIP-seq for protein CHD1, which is a chromatin remodeler and doesn’t have a motif.

Among the top five motifs discovered for the CTCF dataset, the server filtered out three motifs due to their lack of significance (http://bib.umassmed.edu/factorbook/view/fmpCWHiNz).

Fig. 1a shows the significant motif that passed our filtering procedures and it matches the canonical CTCF binding motif. For each motif there is a chart (as the Fig 1a and Fig 1b bottom panels) that shows the significance of the motifs. A real motif should have instances present in many more peak regions compared to flanking regions (as shown in the Fig. 1a, the red line and black line are clearly separated). Also the distance from the motif instance to the summit reveals both the quality of the input peak resolution and the quality of the motif (random motif usually shows larger distances from the summit). Fig. 1b shows a non-significant motif that is filtered out. As explained earlier, the low peak fraction and the similarly low flanking fraction show the randomness of the motif. Also the distance to the summit is larger for the non-significant motif compared to the significant one.

**Fig. 1.**
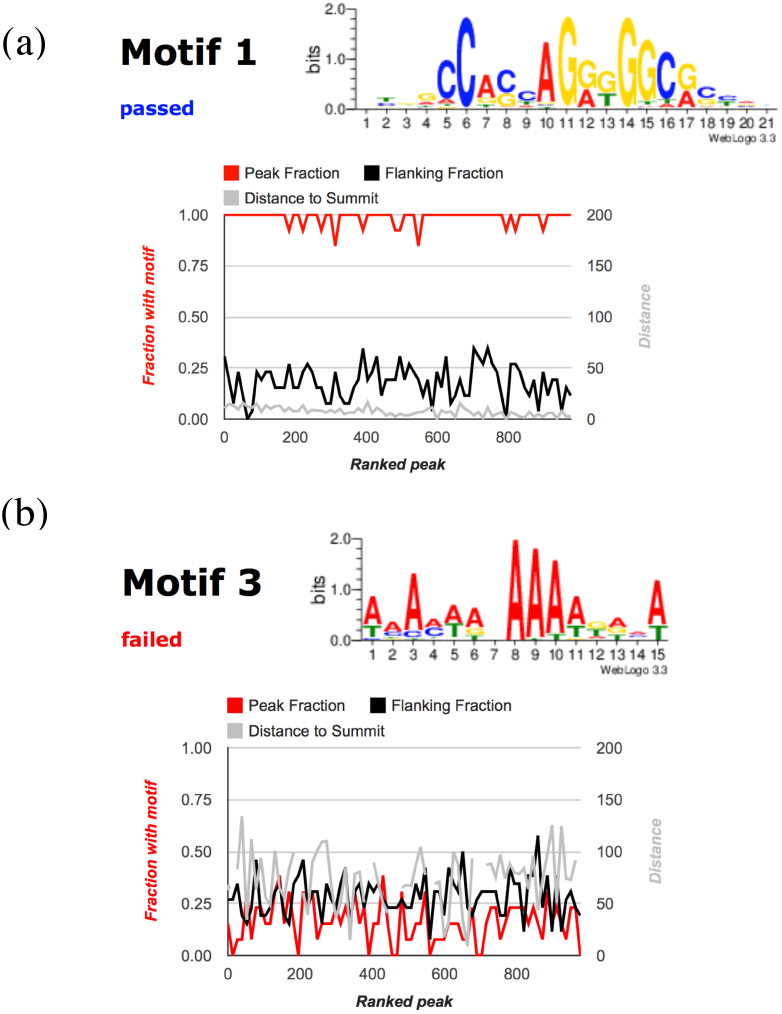
Discovered motifs from *de novo* motif search and their filtering results. (a) shows canonical CTCF motif discovered by MEME, which passed the filtering procedure by random genomic sequences with matching GC contents (*P*-value 0.0) and motif instance enrichment (5.2 folds) in the peak regions compared to the flanking regions. The sequence logo representation of the motif is drawn is shown in the top panel. In the bottom panel, the quality of the motif is represented according to the peak rank or peak intensities from the user input peak file. Peak fraction (red line) represents the fraction of peaks contains the motif instances in each bin and the flanking fraction (black line) represents the fraction of the flanking regions that contains motif instances in each bin. Distance to the summit (gray line) represents the average distance to the summit from the motif instances, (b) shows a random pattern discovered by MEME from the same CTCF dataset, which failed to meet the significance thresholds by random genomic sequences with matching GC contents (*P*-value 1.0) and motif instance low enrichment (0.5 folds) in the peak regions compared to the flanking regions.

For CHD1, Factorbook Motif Pipeline filtered out all discovered motifs reported by MEME as expected. The graphical results are available at the following URL, http://bib.umassmed.edu/factorbook/view/fmplVQ0K0.

## ACKNOWLEDGEMENTS

The authors are grateful to Brian Pierce and Jill Moore for their helpful comments and critical reading of the manuscript.

